# Multimodal measurements of phototoxicity in nonlinear optical microscopy

**DOI:** 10.1101/2021.08.27.457929

**Authors:** Xinyi Zhang, Gabriel Dorlhiac, Markita P. Landry, Aaron Streets

## Abstract

Nonlinear optical imaging modalities, such as two-photon microscopy and stimulated Raman scattering (SRS) microscopy, make use of pulsed-laser excitation with high peak intensity that can perturb the native state of cells. In this study, we investigated the short and long-term effects of pulsed laser induced phototoxicity. We used bulk RNA sequencing, quantitative measurement of cell proliferation, and measurement of the generation of reactive oxygen species (ROS) to assess phototoxic effects, at different time scales, for a range of laser excitation settings relevant to SRS imaging. We define a range of laser excitation settings for which there was no significant ROS generation, differential gene expression, or change in proliferation rates of mouse Neuro2A cells. Changes in proliferation rate and ROS generation were observed under imaging conditions with an excitation intensity of over 600 mW/μm^2^. Repeated imaging of the same field of view at this excitation intensity of over 600 mW/μm^2^ resulted in visual damage to N2A cells. Laser induced perturbations in live cells may impact downstream measurements of cell state including subsequent imaging or molecular measurements. This study provides guidance for imaging parameters that minimize photo-induced perturbations in SRS microscopy to ensure accurate interpretation of experiments with time-lapse imaging or with paired measurements of imaging and sequencing on the same cells.

## Introduction

Advances in optical engineering over the past three to four decades have produced numerous new technologies that have revolutionized the life sciences. Principally among these are the commercial availability of reliable plug-and-play lasers across a broad range of wavelengths, both in the continuous-wave and pulsed regimes, as well as advanced scanning microscopy systems. In response, numerous microscopy techniques have been developed to push the boundaries of what can be probed optically in a biological system. One such set of developments include super-resolution microscopy techniques, such as PALM or STORM^1,2^, which are used to elucidate sub-cellular organization at below the diffraction limit. Perhaps most prominent among these new techniques, however, has been the proliferation of two-photon excited fluorescence (TPEF) microscopy. TPEF microscopy has been adopted widely, particularly in fields such as neuroscience as it has allowed for imaging deeper into tissue due to the wavelengths used for excitation^3^.

Beyond TPEF microscopy, there is a growing field of other nonlinear optical microscopy techniques. Nonlinear optics deals with a regime where the peak optical power becomes large enough that nonlinear effects become important. Such peak power is provided by modern pulsed lasers operating in the picosecond to femtosecond regime which concentrate power into a very short pulse duration. The nonlinear optical microscopy field has garnered a lot interest in the past decade, as the additional modalities it offers are in many cases label-free. In principle this means that samples can be probed without the introduction of additional dyes, and in many cases without fixation. Second harmonic generation (SHG) imaging, for example, has been widely used to investigate structure and order in a number of biological systems such as the cornea and spine, and generally in tissues with a high collagen content^4^. Third harmonic generation (THG) has been used to extend some of the benefits of SHG to centro-symmetric materials^5^.

While SHG/THG provide label-free structural information, and TPEF provide specificity, albeit with a fluorescent label, there has been increasing demand for techniques which provide the advantages of being both label-free as SHG/THG, and specific as TPEF. Confocal Raman imaging fills this niche quite well, as it provides chemical specificity in the form of vibrational signatures; however, traditional spontaneous Raman scattering is an inherently infrequent process requiring long exposure times. As a result, many have turned to the nonlinear varieties of spontaneous Raman scattering, including coherent anti-Stokes Raman scattering (CARS) and stimulated Raman scattering (SRS). These techniques make use of two pulsed lasers of chosen frequency such that the difference in their frequencies is equal to the frequency of a molecular vibration of interest. Such vibrations could, for example, be CH_3_ stretches, which are most abundant in intracellular proteins. SRS in particular has a number of advantages, namely that signal intensity is linear in the concentration of the molecule under investigation, and as such is more easily quantitative, and that an SRS spectrum is identical to a spontaneous Raman spectrum. Both CARS and SRS have been used to investigate numerous systems including infiltrating tumor cells in fresh brain samples^6^ and lipid droplet formation in single cells^7^.Yet these studies have often avoided addressing an important question: what effect does the exposure to pulsed lasers have on the sample under investigation? This question is particularly relevant in cases where the microscopy experiment is not the end-point measurement, as in cases where imaging is paired with, for example, sequencing.

Phototoxicity is often mentioned in the context of microscopy, but is rarely thoroughly investigated, and often less so in the pulsed laser regime. Even in the continuous-wave regime, visible laser radiation has been associated with increase in reactive oxygen species (ROS) and loss of cell viability^8,9^. Imaging with pulsed lasers has also been associated with a decrease of cloning efficiency^10^ and membrane blebbing,^11^ in addition to an increase in ROS and loss of cell viability^12–14^. Yet despite these observations, and widespread acknowledgment that phototoxicity is an issue, there lacks a comprehensive study of how laser irradiation alters cell state, on the metabolic and transcriptomic level.

Given the promise of nonlinear imaging techniques, and particularly label-free ones such as SRS, this seems to be an important uncertainty to address. This is particularly the case as interest continues to grow for paired measurements, where samples are imaged prior to an additional measurement. Paired imaging and sequencing experiments on the same samples, for example, have led to numerous insights. Lane *et al*. studied the relation between the dynamics of nuclear factor κB activation and gene expression by paired fluorescent imaging and single-cell RNA-seq^15^. Recent advances provided high-throughput methods to combine RNA-seq and microscopy on the same cells, such as μCB-seq^16^ and SCOPE-Seq^17^. To accurately interpret the paired RNA-seq data given the reported phototoxic effects of laser-based imaging, it is important to understand the impact of laser radiation on the transcriptome.

To address this need, this study aims to uncover the cellular response to pulsed laser excitation. Particularly, we investigate the effects of stimulated Raman scattering microscopy (SRS), which as a multi-pulse technique, often reaches some of the highest photon fluxes commonly used. We used two pulsed excitation pulses, as is common in SRS, and probed the effects of single pulses, relevant to all major nonlinear imaging techniques, as well as two-pulse excitation when the pulses are not temporally coincident. Measurements of reactive oxygen species generation, induced by laser excitation, along with quantification of cell proliferation after exposure, were used to compare to previous studies^10,12^. Critically, we tested for potential phototoxic effects in gene expression using RNA-sequencing, which allows for a comprehensive, transcriptome-wide assessment of potential changes caused by laser exposure. The results of this study provide comprehensive guidelines for minimizing the effects of photo-toxicity in SRS imaging, and nonlinear imaging in general.

## Methods

### 1. Cell culture

Neuro2A cells (N2A; UCB Cell Culture Facility) were cultured in T25 flasks using standard cell culture conditions for all experiments - 37 °C, 5% CO_2_, and complete growth medium consisting of DMEM (Gibco^™^ 10566016), 10% FBS (Avantor Seradigm 89510), and 1% penicillin-streptomycin (Gibco^™^ 15140122). Cells were grown to approximately 80-90% confluency and one day prior to each imaging experiment, they were resuspended and seeded into 8-well glass-bottom μ-Slide (ibidi 80827) for the ROS assays and into 384 glass-bottom well plates (Corning 4581) for the proliferation assays and RNA-seq. The growth medium for the ROS assays did not contain phenol red. At the start of each experiment, the confluency was approximately 40% for the proliferation assays and 70% for the ROS assays and RNA-seq.

### 2. Live-cell imaging and pulsed laser exposure

The optical setup has been described previously^18^. The fundamental and tunable output from a commercially available femtosecond oscillator/OPO (Insight DS+, Spectra-Physics) was used as the excitation source. The OPO output was tuned to 796 nm for all measurements. The fundamental was at 1040 nm. For single-pulse measurements, only the OPO output at 796 nm was used. For two-pulse measurements the output from both the fundamental and OPO were used. Each line’s power was controlled through the use of a variable attenuator consisting of a half-wave plate or half-wave fresnel rhomb (OPO output), and a polarizer, and set to the power specified in the main text. The inter-pulse delay was controlled by a delay stage (FCL200, Newport) on the tunable output line, and the two lines were combined on a 1000 nm short-pass dichroic mirror (Thorlabs), and fed into an inverted scanning microscope. (Olympus IX83-FV1200) A 20x objective with a numerical aperture of 0.75 (Olympus, UPLSAPO20X) was used for all measurements, and frames were acquired at 512 x 512 pixels with a dwell time of 2 μs per pixel. The wavelength of 796 nm was chosen because when coincident with the fundamental 1040 nm line it provides the pump for a stimulated Raman interaction at a frequency of 2950 cm^-1^, corresponding to CH3 stretches, largely found in proteins throughout the cell. In addition to SRS imaging, to deconvolve the contribution of the stimulated Raman process from the other effects of the dual laser excitation, measurements were acquired at an inter-pulse delay of 50 ps, with the 796 nm pulse arriving first.

### 2. ROS assay

The N2A cells were exposed to the different settings as described in Table 1, with three replicates per exposure condition. The three negative control wells were not exposed. ROS generation was detected by CellROX^™^ Green reagent (Invitrogen) following the manufacturer’s protocol. The dye was diluted to 5 μM in pre-warmed complete growth medium immediately before addition to cell culture. The growth medium in wells for exposure and the negative control wells was replaced by the medium with the dye in each well after the laser exposure for that well was completed, followed by 30 min incubation. After incubation, each well was washed three times with warm medium. A fluorescent image was taken in the GFP channel after the wash. A bright-field image of the same field of view was taken for each fluorescent image, so that the fluorescent increases post-exposure were normalized by the number of cells in each field of view. The same protocol was followed in empty wells to confirm that the laser beams did not react with CellROX Green dyes directly to cause any fluorescent changes. Cell boundaries were manually defined using the bright-field images to measure fluorescence post-exposure. The fluorescence was normalized to the background by manually selecting regions in the same well that did not have cells. A flat-field correction was also applied to each fluorscence image. The reference for the flat-field correction was taken in empty wells with the same concentration of CellROX Green in warm complete medium under the same settings for the GFP channel. During all procedures, N2A cells were kept at 37 °C with 5% CO_2_.

**Table 1.**
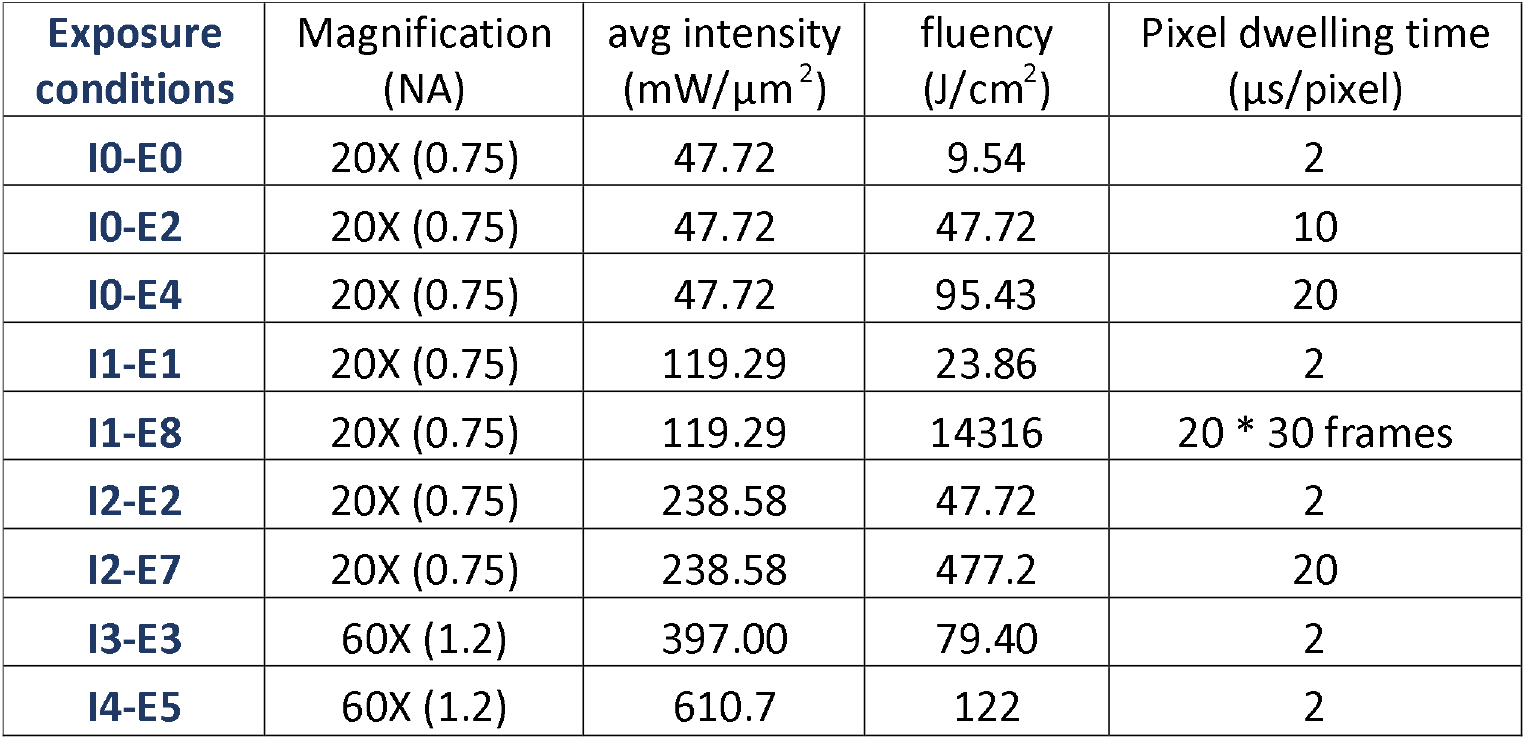
Summary of laser exposure conditions tested in this study.

### 3. Proliferation assay

For each condition listed in Table 1, the entire culture wells in a 384 glass-bottom well plate were exposed to laser excitation by dividing each well into multiple non-overlapping fields of view. The stage was moved during the imaging of a well to scan each field of view once with the laser, such that all the cells in the same well were subjected to the same laser exposure condition. On the same 384-well plates, three wells of negative control cells were not exposed to the laser but were otherwise subjected to the same conditions. To measure the proliferation overnight with time-lapse imaging, we used IncuCyte ZOOM ((Essen BioScience) with a 10X objective to automatically acquire phase-contrast images while keeping the 384-well plates at 37 °C and 5% CO_2_. A phase-contrast image was taken every 30 min to 2 hours inside the IncuCyte for each well. N2A cells in the images were segmented using an algorithm for segmenting cells in bright-field images from Buggenthin *et al* ^19^. Fiji was used for counting cells in the binary images after segmentation^20^.

### 4. RNA-seq

As in the proliferation assays, for each RNA-seq condition, the entire cell culture well in 384 glassbottom well plates was exposed to laser excitation, such that all the cells in the same well were subject to the same laser exposure. The cells were lysed 1 hour after laser exposure to allow sufficient time for transcriptional changes^21^. Three wells of unexposed cells subjected to the same conditions were lysed at the same time. We followed the manufacturer’s protocol for RNA extraction (Qiagen RNeasy Mini Kit) and library preparation (NEBNext Ultra II RNA library prep kits for Illumina). Differential gene expression was tested for each exposure settings against the negative controls that were subject to the same ambient conditions and the same library preparation protocols but with no laser excitation. A false discovery rate < 0.05 was used as the threshold for differential expression after adjusting for multiple hypothesis testing with the Benjamini-Hochberg procedure^22^ or with the Bonferroni correction^23^. The R package limma was used to perform this differential expression analysis.^24^

## Results

### 1. Mechanism of Laser-induced Damage

Photodamage induced by multiphoton microscopy includes both linear and nonlinear processes^11,25,26^. It has been suggested that, with lower laser peak power in CARS imaging, photodamage is dominated by linear dependence on peak power^11^. Whereas second order processes dominate at higher peak power^11^. The linear dependence of photodamage on peak power has been associated with one-photon absorption in human skin and *Escherichia coli*^11,27,28^. Linear and higher order processes can both lead to heating, but result in different profiles of temperature distribution where nonlinear absorption is mediated by free electrons^29^. Plasma generation through ionization, a result of the multiphoton process, can also have chemical effects in addition to thermal. The chemical effects include increase of ROS generation and fragmentation of biomolecules, e.g. DNA fragmentation and loss of membrane integrity^8,12,13,29^. ROS-mediated phototoxic effects were associated with thermal inactivation of ROS scavengers in cells, in addition to the increase in ROS generation^8^. In the same study, PCR arrays, immunoblotting, and ATF-knockdown were used to assess the role of ER stress pathway after laser irradiation^8^. Using a continuous wave laser to achieve a fluency comparable to our study (around 27 J/cm^2^), Khan *et al*. only observed phototoxic effects in black wells that absorbed 100% laser irradiation but not in clear wells^8^. Our study used clear glass-bottom well plates to minimize the amount of laser irradiation absorbed by the well plates. Damage in DNA and the plasma membrane has been reported to result in apoptosis^12^. However, some studies found increase in ROS generation or apoptosis in cells exposed to laser without direct DNA damage^8,13^. Other observed cellular damages that did not directly lead to apoptosis include non-lethal morphological changes, loss of cloning efficiency and uncontrolled cell growth^10,14,25^. Understanding the thresholds for these phototoxic effects is critical for designing a stimulated Raman scattering imaging experiment. We aimed to measure the transcriptome-wide response in N2A cells by performing bulk RNA-sequencing after exposure to femtosecond pulsed laser irradiation with different exposure settings. We also investigated both the immediate and long-time scale response by quantifying intracellular generation of ROS immediately after irradiation and by characterizing cell proliferation for 24 hours after exposure.

### 2. Laser induced phototoxicity with typical SRS imaging conditions

In stimulated Raman scattering (SRS) microscopy, as well as two-photon excitation microscopy (TPEF), many critical imaging parameters can influence the degree of photodamage on cells, including laser peak intensity, average intensity, repetition rate, excitation wavelength, total energy deposition on the samples, and Raman resonance^10,11,30^. We tested the effects of laser exposure on Neuro2A (N2A) cells, a mouse neuroblastoma cell line to investigate how the choice of parameters contributes to photodamage. The imaging settings were chosen to represent typical settings of SRS imaging of proteins and were also comparable to settings used for TPEF.

First, we tested the long-term effect of the average excitation intensity on N2A cells in two sets of imaging experiments with identical cells, by tuning laser excitation power. The repetition rate and pulse width were kept constant at 80 MHz and 120 fs. Thus, the test was equivalent to comparing different laser peak intensities. N2A cells were exposed to a single 120-fs laser with an average power at the sample plane of 20mW and 100mW, corresponding to average intensities of 47.72 mW/μm^2^ (exposure condition I0-E0) and 238.58 mW/μm^2^ (exposure condition I2-E2), respectively at 796 nm. After exposure, phototoxicity was assessed by measuring the proliferation rates of the N2A cells in each well. Additionally, negative control (NC) experiments were performed by seeding cells from the same passage in adjacent wells that did not receive laser irradiation. Proliferation of these negative control cells was measured in parallel with the two imaging conditions. For all conditions and controls, total cell counts were recorded every 2 hours for 30 hours starting about 10 to 20 minutes after exposure. Proliferation was calculated as the fold change in total cell count with respect to the initial counts of cells at time zero. The proliferation rates of the two exposure conditions, I0-E0 and I2-E2, were identical to the negative control group during the 30 hours after laser exposure (Figure 1a), indicating that these intensity levels did not have adverse effect on the cloning efficiency of N2A cells.

**Figure 1.**
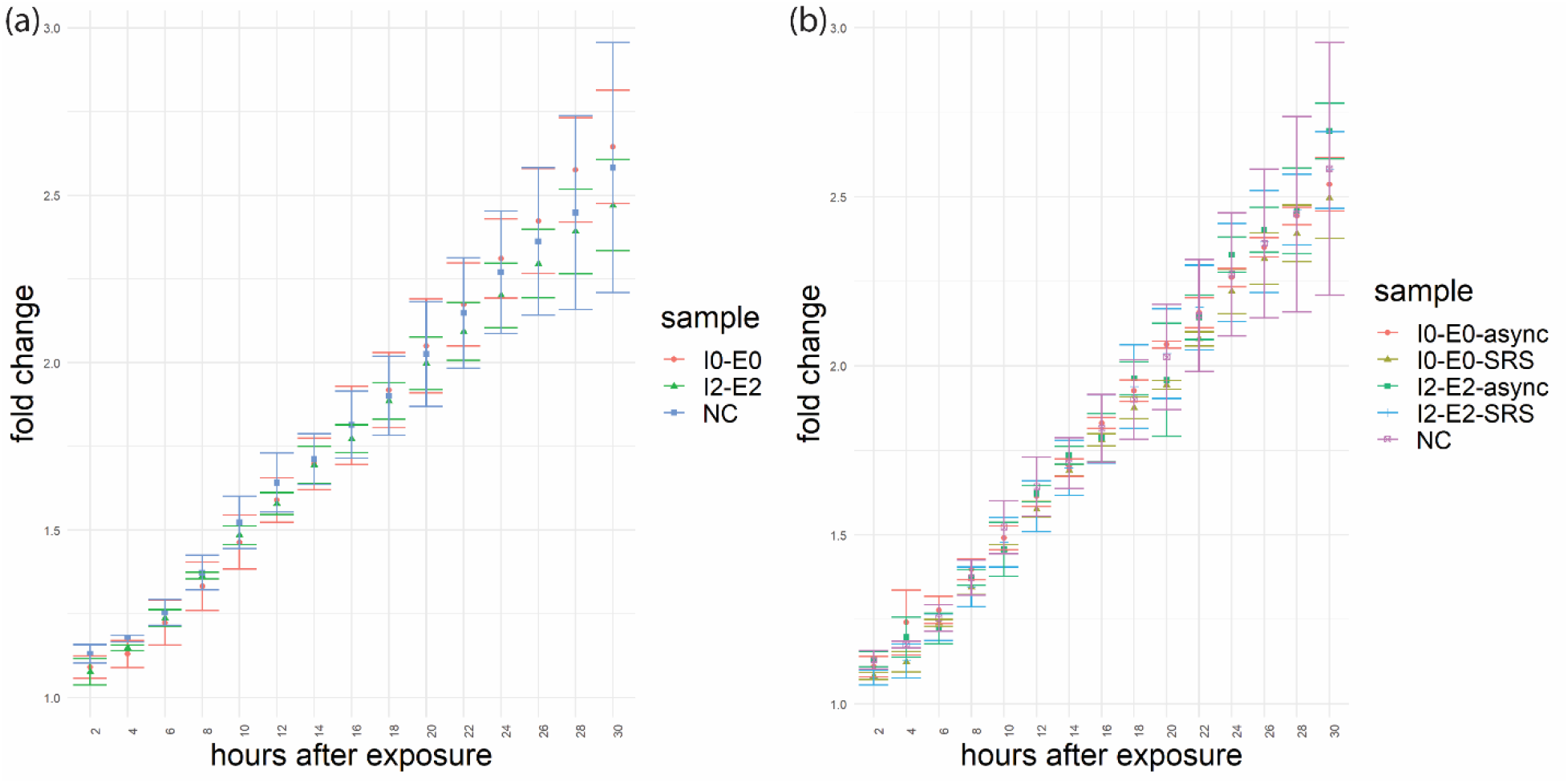
Proliferation rates after laser exposure. Measured as fold changes of the counts of N2A cells every 2 hours with respect to the initial counts of cells immediately after the completion of all laser exposures. I0: average intensity = 47.72 mW/μm^2^; I2: average intensity = 238.58 mW/μm^2^. E0: fluency = 9.54 J/cm^2^; E2: fluency = 47.72 J/cm^2^; NC = negative control; SRS: cells were imaged with SRS; async: cells were imaged with an interpulse delay of 50 ps. a) A single pulsed laser at 796 nm was used. b) Two lasers at 796 nm and 1040 nm were used, which had 1:1 power ratio. The total average intensities of the two laser settings were the same as the respective single laser settings.

In addition to the photo-damage caused by the combined power from two excitation sources, the coherent Raman scattering process can also induce photodamage caused by Raman resonance^11^. We tested if excitation with pump and Stoke’s wavelengths chosen to excite Raman resonance in CH_3_ bonds would result in damage that perturbed proliferation rates. To do this, we added a second excitation source and chose the intensity of each source to achieve a total combined intensity that was equivalent to the previous settings. For each of the two intensities, 47.72 mW/μm^2^ and 238.58 mW/μm^2^, N2A cells were imaged with SRS by dividing the total intensities between two lasers at 796 nm and 1040 nm with 1:1 ratio (exposure condition I0-E0-SRS for 47.72 mW/μm^2^ and I2-E2-SRS for 238.58 mW/μm^2^). We also tested whether any phototoxic effect from the Raman resonance was offset by allocating half the total intensity to a longer wavelength. We exposed separate samples of N2A culture to the same two-laser settings but eliminated the Raman resonance by delaying the arrival time of the 1040 nm pulse by 50 ps with respect to the 796 nm pulse (exposure condition I0-E0-async for 47.72 mW/μm^2^ and I2-E2-async for 238.58 mW/μm^2^). We observed that neither condition diminished cell proliferation over the 30-hour period immediately after laser exposure, comparing to the negative control cells that were not exposed to laser irradiation (Figure 1b).

The lack of change in N2A proliferation rate indicates if there was any perturbation caused by the laser exposure under these settings, this perturbation did not negatively impact the long-term survival of N2A cells. This, however, does not preclude the possibility of shorter timescale photo-induced perturbations that do not affect proliferation rate. Such shorter timescale perturbations might be recorded in transcriptional activity which occurs on timescales that are much shorter than the doubling time. To test for the potential changes in transcription, we performed bulk RNA sequencing (RNA-seq) on N2A cells that were exposed to the single-laser excitation settings tested in the proliferation assays (exposure condition I0-E0 and exposure condition I2-E2). For each sample exposed to laser irradiation, RNA was extracted 1 hour after exposure. The length of post-exposure incubation was chosen to allow sufficient time for transcriptional changes to occur before significant mRNA degradation, according to studies of mammalian cell transcription rate^31^ and mRNA half-life^32^. We compared gene expression in the exposed cells to negative control samples that were not exposed to laser irradiation but were otherwise subjected to the same handling. In this differential expression analysis, we did not find any differentially expressed genes (DEGs) with a false discovery rate (FDR) < 0.05, after adjusting for multiple hypothesis testing with the Benjamini-Hochberg procedure^22^ or with the Bonferroni correction^23^ (Figure 2a, 2b), indicating that these excitation intensities produced no significant perturbation to gene expression in N2A cells.

**Figure 2.**
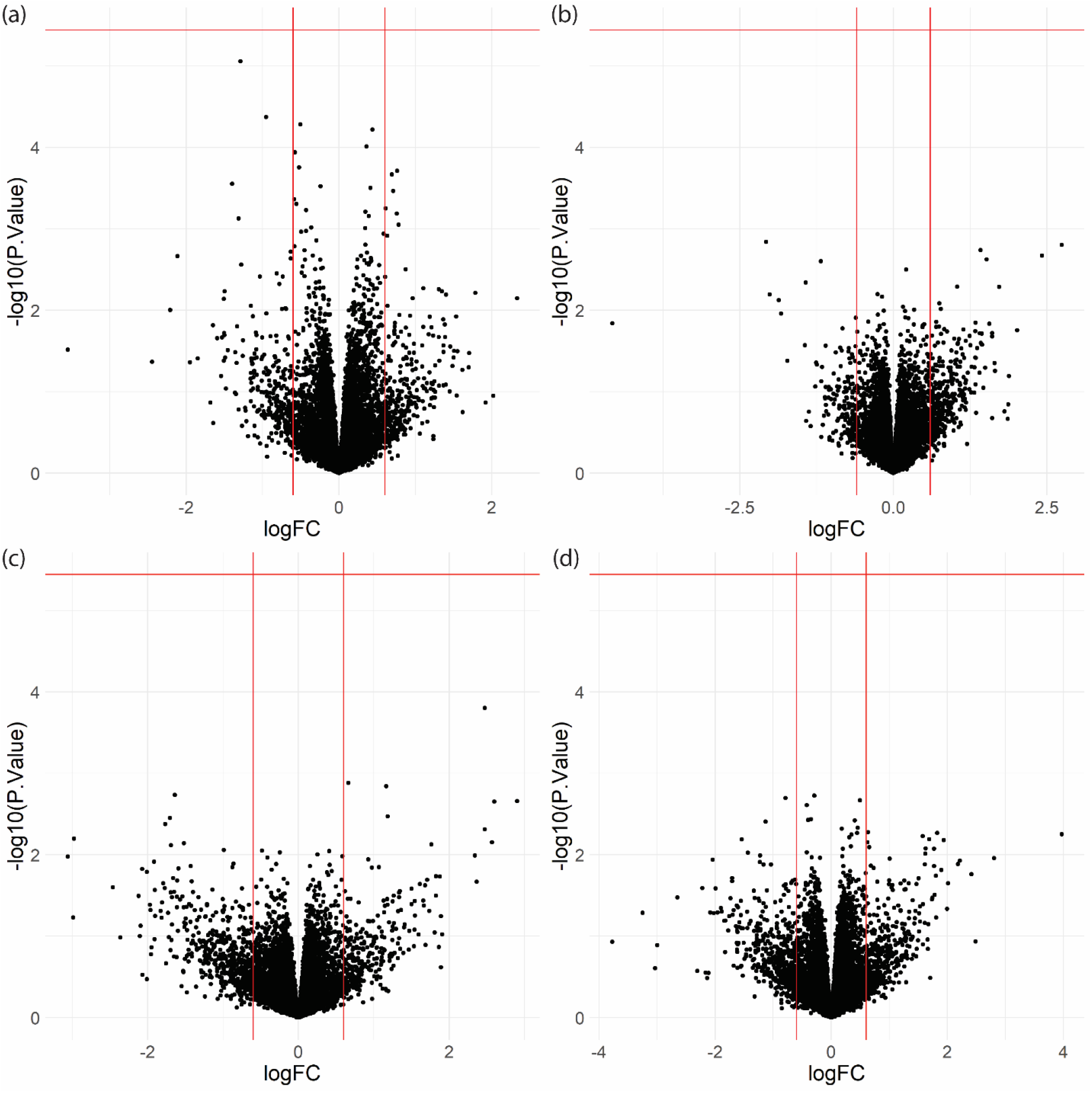
RNA-seq volcano plots of differential gene expression from single-laser exposure conditions compared to the negative control samples. a) I0-E0: average intensity = 47.72 mW/μm^2^; fluency = 9.54 mW/μm^2^. b) I2-E2: average intensity = 238.58 mW/μm^2^, fluency = 47.72 J/cm^2^. c) I0-E2: average intensity = 47.72 mW/μm^2^; fluency = 47.72 J/cm^2^. d) I0-E4: average intensity = 47.72 mW/μm^2^, fluency = 95.43 J/cm^2^. logFC: log2 fold change with respect to negative control samples (NC) that were not exposed to laser. Vertical lines are at −0.6 and 0.6. −log10(p-value): −log10 of p-values of differential expression with respect to NC. Horizontal lines indicate familywise error rate of 0.05 after Bonferroni correction^23^.

In addition to laser excitation intensity and Raman resonance, the total energy deposition on the cells could also influence the extent of phototoxicity. This incident photon dose is determined by the excitation intensity and the total exposure time. In the context of laser scanning microscopy, slower scanning speeds or repeated imaging can increase photo-induced damage^11,30^. We tested the impact of increasing energy deposition per unit area (fluency) by increasing pixel-dwelling time given a fixed intensity. To achieve an increase in fluency of 5x and 10x, the pixel dwelling times were increased from 2 μs/pixel in the 47.72 mW/μm^2^ samples (I0-E0) to 10 μs/pixel and 20 μs/pixel (exposure condition I0-E2 and I0-E4) respectively. We performed RNA-seq with the same procedure as in the previous test on these two groups with higher fluency. Comparing to the negative control groups that did not undergo laser exposure, no significant differential gene expression was found in these two exposure groups either (Figure 2c, 2d).

### 3. Laser-induced photo-toxicity with high-intensity SRS imaging

We did not observe any phototoxic effects on proliferation or gene expression with the typical SRS imaging parameters. However, in signal-limited SRS imaging applications much higher excitation intensities are often implemented, such as the identification of the signature of neuronal membrane potentials^33^. In these scenarios, researchers must balance the large excitation intensities required for high-sensitivity SRS with photo-damage in the sample resulting from strong photo-absorption including plasma generation^11^ and extreme heating^34^. This damage can often be observed visually as burning or boiling of the sample^35^. Thus, we probed the impact of laser absorption at settings that are close to causing sample destruction by heating. We increased the total average intensity to 397 mW/μm^2^ by switching from a 20X to 60X objective with 1.2 NA. We chose a near-burning condition by repeatedly imaging the same field of view until burning was observed in some cells (Figure S1), which occurred after 14 successive image scans, the equivalent of 1111.6 J/cm^2^. In this setup, two lasers at 796 nm and 1040 nm with a 10:3 power ratio and 2 μs pixel dwelling time were used. In the subsequent experiments, we used the same setup but only scanned each field of view once (exposure condition I3-E3-SRS, Table 1). To test for the long-term survival, we performed a proliferation assay following the same procedure as in the previous section. No change was observed in the proliferation rate of N2A cells after exposure to this near burning condition using two lasers with pulses that were not temporally coincident compared to the negative control group without laser exposure (Figure 3b). Switching to imaging with SRS where the two lasers with identical settings were temporally coincident, allowing for Raman resonance at 2950 cm^-1^, also did not change the proliferation rate of N2A cells (Figure 3a). We investigated whether these imaging conditions caused shorter time-scale perturbations reflected as transcriptional changes by performing RNA-seq 1 hour after imaging the N2A cells with SRS under the near-burning condition (I3-E3-SRS), as was described in the previous section. No differentially expressed gene with a FDR < 0.05 were found in this condition compared the negative control group (Figure 3c). This result suggests that N2A cells are robust to high-intensity laser exposure and vibrational excitation.

**Figure 3.**
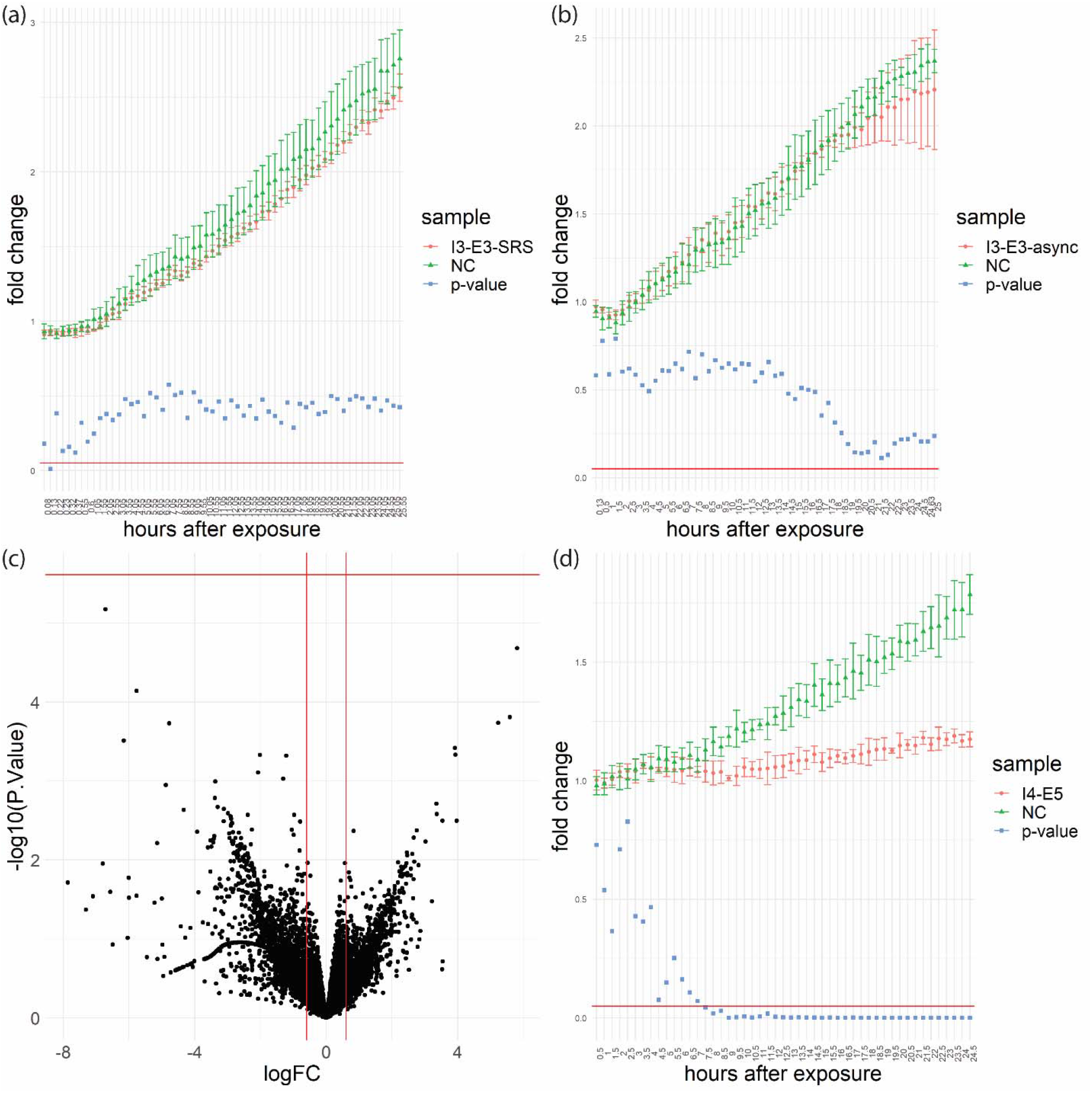
Proliferation and RNA-seq of near-burning conditions. (a) Proliferation rates of I3-E3-SRS measured as fold changes of the counts of N2A cells at every 30 minutes with respect to the initial counts of cells immediately after the completion of all laser exposures for 25.5 hours; (b) proliferation rates of I3-E3-async which has the same laser settings as in (a) but has laser pulses that arrive at the sample at different times (50 ps interpulse delay); (c) Volcano plot of RNA-seq results of cells exposed to I3-E3-SRS with 1 hour post-exposure incubation; (d) proliferation rates of I4-E5 measured at every 30 minutes for 24.5 hours after laser exposure. I3: average intensity = 397 mW/μm^2^; I4: average intensity = 610.7 mW/μm^2^; E3: fluency = 79.4 J/cm^2^; E5: fluency =122 J/cm^2^; NC = negative control. SRS: cells were imaged with SRS; async: cells were imaged with an interpulse delay of 50 ps. logFC: log2 fold change with respect to negative control samples that were not exposed to laser. Vertical lines are at −0.6 and 0.6. −log10(p-value): −log10 of p-values of differential expression with respect to NC. Horizontal lines indicate familywise error rate of 0.05 after Bonferroni correction^23^.

We increased the intensity further to probe the limits of non-perturbative femtosecond laser scanning excitation. With an average intensity of 610.7 mW/μm^2^ at 796 nm (exposure condition I4-E5) and 2 μs pixel dwelling time, we observed cell burning starting around the second or third sequential imaging frame (equivalent to 244 to 366 J/cm^2^). Under the same laser excitation condition but only one imaging frame, the samples exposed to laser irradiation had a proliferation rate that was significantly reduced (~25%) compared to the negative control samples that received the same treatment except laser exposure (Figure 3d). Laser-induced changes in N2A started to result in proliferation rate changes between average intensity of 397 mW/μm^2^ with a fluency of 79.4 J/cm^2^ and 610.7 mW/μm^2^ with a fluency of 122 J/cm^2^.

### 4. Photo-induced generation of reactive oxygen species in N2A cells

Generation of various reactive oxygen species (ROS) has been observed under a wide range of laser exposure conditions. The generation of ROS has been associated with processes related to sample ionization^11^, heat generation^8^, and active repair mechanism of cells^13^. Measurement of the generation of ROS is an additional approach for the investigation of photo-induced physiological perturbations in live cells, that occur on shorter timescales and that may not have observable influence on gene expression or proliferation rate. Although the causality between ROS generation and change in gene expression is unclear, such measurements can be used as an orthogonal measurement of photo-induced perturbations in live cells. We explored a series of increasing excitation intensities and pixel dwelling times that were used in the RNA-seq and proliferation assays as listed in Table 1. The increase in ROS generation was measured by the increase in fluorescence of the CellROX Green dye, which became fluorescent after oxidation by ROS (Figure 4a). Comparing to the negative control samples without laser exposure (Figure 4g, 4h), the only condition that resulted in a significant ROS increase was I4-E5 (average intensity = 610.7 mW/μm^2^, fluency = 122 J/cm^2^; Figure 4c, 4d), the near-burning condition we described in the previous section that also resulted in decreased proliferation rates. This is also the exposure condition with the highest intensity among all experiments. All other exposure settings tested did not result in significant generation of ROS measured by the CellROX Green dye. This includes the same exposure settings that did not result in changes of proliferation rates or differential gene expression measured by RNA-seq in the previous tests (exposure condition I0-E4, I2-E2, I3-E3; Figure 4b, 4e, 4f). This observation confirms that these imaging conditions caused insignificant perturbations on the N2A cells. Additionally, we tested two conditions that had lower intensity but higher fluency than I4-E5, the condition with ROS increase. First, we used the same intensity as I2-E2, 238.58 mW/μm^2^, using 796 nm wavelength and 20X objective. The fluency was increased to 477.2 J/cm^2^ by increasing the pixel dwelling time to 20 μs. This condition with 3.9 times higher fluency than I4-E5 did not result in significant ROS generation. We further increased the fluency to 14316 J/cm^2^ using an average intensity of 119.29 mW/μm^2^ at 796 nm. The fluency was achieved by using a 20 μs pixel dwelling time and repeatedly imaging the same field of view 30 times. This 117 times increase in fluency from I4-E5 also did not cause in increase of ROS generation. Thus, intensity rather than fluency was likely to be the major factor determining whether N2A cells had an increase in ROS generation after exposure to a particular laser excitation condition. If high imaging intensity is necessary for a particular application, the addition of ROS scavengers, such as N-Acetyl-L-cysteine and Catalase, might protect cells from oxidative damage^8^. Lowering cell culture temperature during imaging has also been reported to reduce photodamage caused by exposure to continuous-wave lasers^8^. Additional experiments are needed to examine if the addition of oxygen scavengers or lowering temperature would result in changes of gene expression or other perturbations.

**Figure 4.**
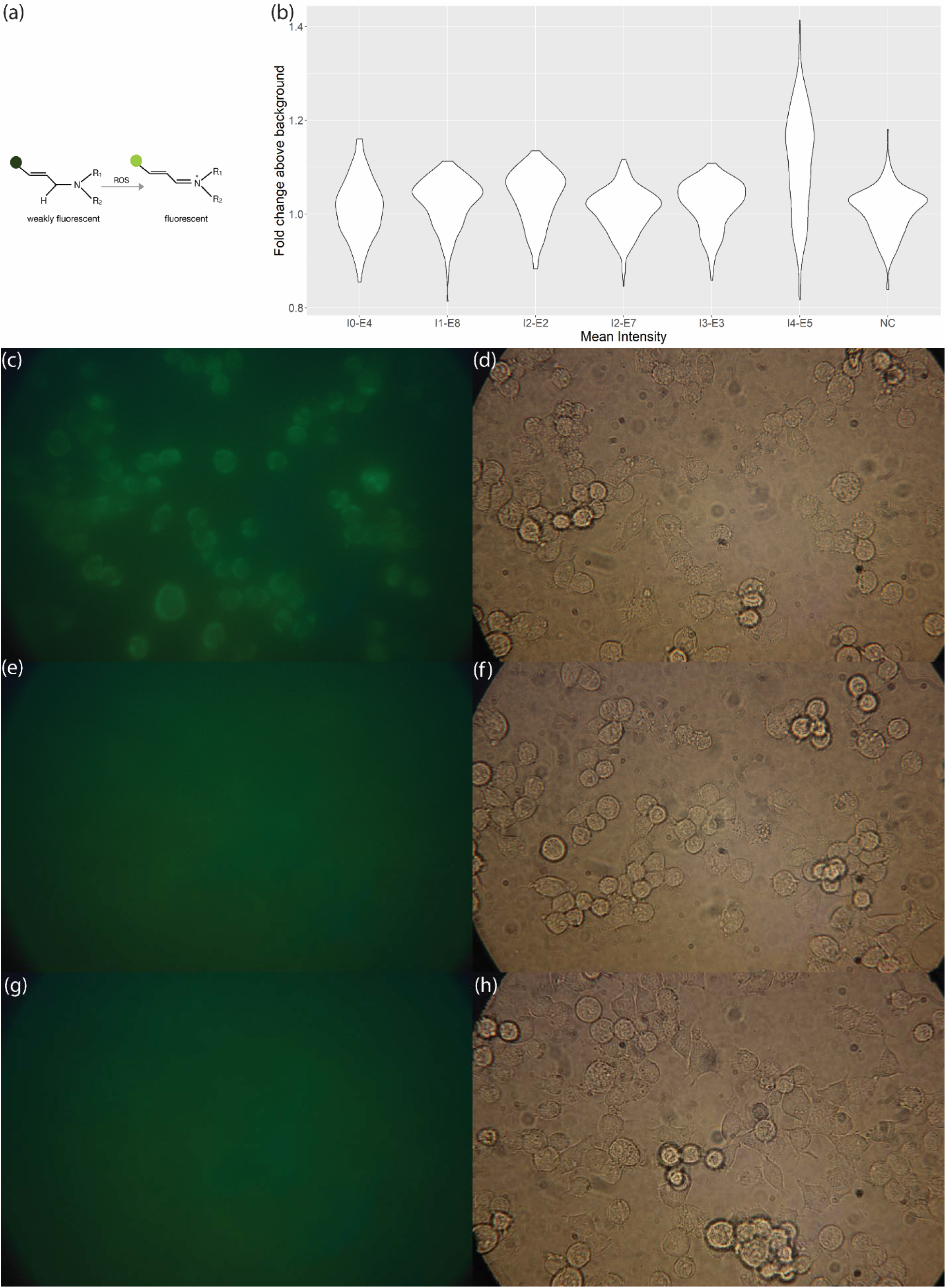
ROS measurements across laser intensities and fluencies. (a) Skeletal formula schematic of oxidation of CellROX Green dye by ROS^36^ (b) Distribution of mean intensities of all cells in samples with laser exposure and the negative control group after flatfield correction. p-value of I0-E4 vs NC is 0.0871. p-value of I1-E8 vs NC is 1.47e-5. p-value of I2-E2 vs NC is 2.89e-10. p-value of I2-E7 vs NC is 0.33. p-value of I3-E3 vs NC is 9.11e-4. p-value of I4-E5 vs NC is 5.73e-45. (c) GFP channel after exposure to I4-E5 and 30 min post-exposure incubation (d) Bright-field image of the same field of view as (c) (e) GFP channel after exposure to I3-E3 and 30 min post-exposure incubation (f) Bright-field image of the same field of view as (e) (g) GFP channel after 30 min incubation of unexposed cells (h) Bright-field image of the same field of view as (g) I0: average intensity = 47.72 mW/μm^2^; I1: average intensity = 119.29 mW/μm^2^; I2: average intensity = 238.58 mW/μm^2^; I3: average intensity = 397 mW/μm^2^; I4: average intensity = 610.7 mW/μm^2^; E2: fluency = 47.72 J/cm^2^; E3: fluency = 79.4 J/cm^2^; E4: fluency = 95.43 J/cm^2^; E5: fluency = 122 J/cm^2^; E7: fluency = 477.2 J/cm^2^; E8: fluency = 14316 J/cm^2^; NC = negative control.

## Conclusion

This study assessed the effects of laser exposure on mouse Neuro2A cells during SRS imaging or other femtosecond pulsed laser-based imaging modalities that use similar excitation parameters. Multiple types of measurements were used – RNA-seq, proliferation assay, and ROS assay – to probe N2A cells at a range of time points after laser exposure. We demonstrated that N2A cells can have strong tolerance to laser exposure over a wide range of imaging power and intensity. This tolerance of N2A cells makes it possible to perform time-lapse imaging, RNA-seq, and other measurements after the first exposure to laser excitation without inducing biological perturbations in the downstream measurements. This allows for accurate interpretation of paired imaging and RNA-seq measurements, which could gain additional insights compared to the individual measurements alone. However, any RNA-seq measurements could only capture the state of gene expression at the time of the measurement. Although we chose the 1-hour incubation time before RNA-seq to allow for sufficient time for potential transcriptomic changes to occur, gene expression changes might happen at a longer time scale, that was not captured in this study. The lack of change in the 1-hour period after laser irradiation could potentially be indicative of the absence of other forms of cellular changes. However, it’s possible that the response of N2A to laser exposure was reflected in changes unrelated to gene expression differences. This study focused on one particular cell type, N2A, and the response might differ from other cell types in their sensitivity and tolerance to laser exposure. The ROS assays also demonstrated that cells can have different tolerance to increase in laser excitation intensity and fluency, which could be a factor to consider when higher resolution is required. Except for I1-E8, where we repeatedly imaged N2A cells in the same field of view 30 times before the ROS assay, we did not explore the possibility of photo-induced perturbations caused by repeated exposure in other contexts. Changes in N2A cells might be observed with extended exposure beyond 20 μs/pixel and 1 imaging frame prior to RNA-seq or proliferation assay. The tunable parameters in SRS imaging form a multidimensional space that allow the imaging settings to vary widely among different applications. The subset of parameters we chose to assess aimed to cover representative changes in intensity, pixel dwelling time, wavelength, and vibrational resonance. Other studies of photodamage in the context of CARS and TPEF observed perturbations to cells using pico-second lasers and/or higher pixel dwelling time (e.g. 60 μs/pixel comparing to 2-20 μs/pixel in our study) ^10–12^. These studies usually examined lower average intensity but higher fluency than our imaging condition with the highest average laser intensity in this study (610.7 mW/μm^2^). Additionally, depending on the timing of downstream measurements, the impacts of laser excitation could manifest in different aspects of cellular processes. Our multimodal measurements at representative timescales after laser exposure could provide insights for assessing potential impacts of laser exposure in future SRS imaging experiments.

## Contributions

A.S. and X.Z. designed the study. X.Z. and G.D. performed experiments. G.D. designed and assembled the SRS microscope. X.Z. analyzed data. M.L. and A.S. supervised the research.

## Acknowledgements

The authors thank Dr. Mary West for the help in the IncuCyte operation and UC Berkeley High-Throughput Screening Facility for the access to the IncuCyte. X.Z. was supported by the UC Berkeley Haas Scholarship. This material is based upon work supported by the National Science Foundation under Grant No. 1845623. A.S. and M.L. are Chan Zuckerberg Biohub Investigators. A.S. is a Pew Scholar in the Biomedical Sciences, supported by the Pew Charitable Trusts. M.L. is supported by the NIGMS of the National Institutes of Health under award number R35GM128922.

## Supplemental Figures

**Figure S1.**
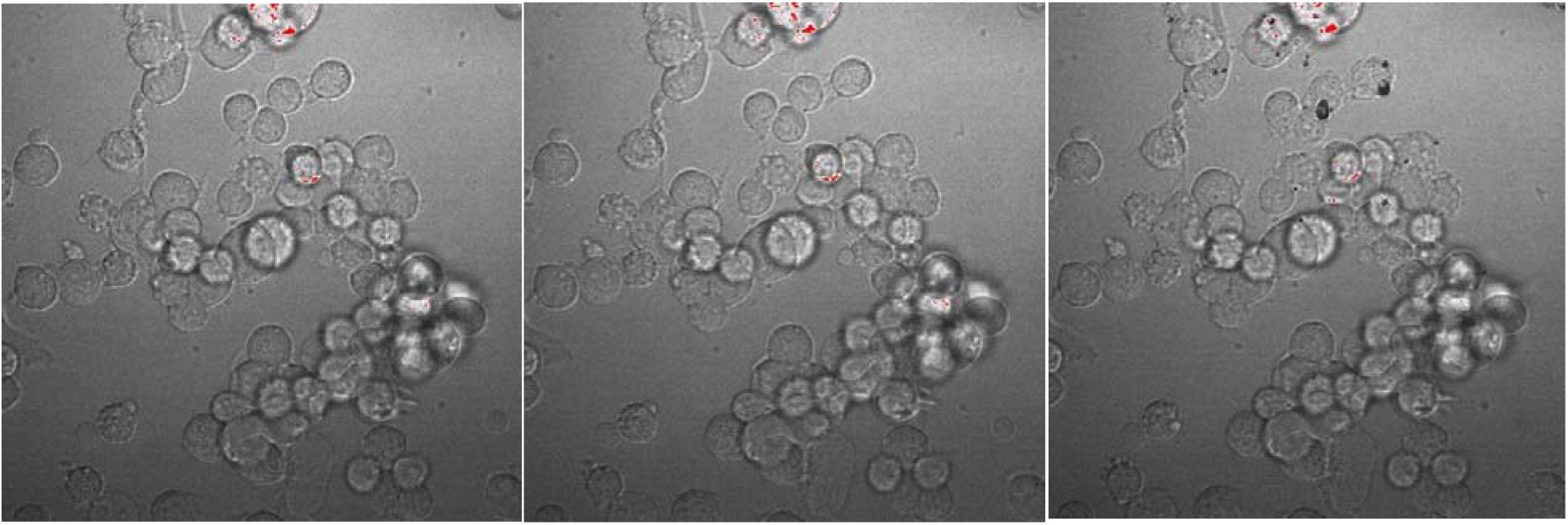
2^nd^, 14^th^, and 51^st^ frames of N2A cells exposed to near-burning condition (I3-E3). I3: average intensity = 397 mW/μm^2^; E3: fluency = 79.4 J/cm^2^.

## Notes

### Competing Interest Statement

The authors have declared no competing interest.

